# Complex evolutionary history of the Y chromosome in flies of the *Drosophila obscura* species group

**DOI:** 10.1101/848804

**Authors:** Ryan Bracewell, Doris Bachtrog

**Affiliations:** Department of Integrative Biology, University of California Berkeley, Berkeley, CA 94720, USA

## Abstract

The *Drosophila obscura* species group shows dramatic variation in karyotype, including transitions among sex chromosomes. Members of the *affinis* and *pseudoobscura* subgroups contain a neo-X chromosome (a fusion of the X with an autosome), and it was shown that ancestral Y genes of Drosophila have become autosomal in species that contain the neo-X. Detailed analysis in species of the *pseudoobscura* subgroup revealed a translocation of ancestral Y genes to the small dot chromosome of that group. Here, we show that the Y-dot translocation is restricted to the *pseudoobscura* subgroup, and translocation of Y genes in the *affinis* subgroup followed a different route. We find that most ancestral Y genes moved independently to autosomal or X-linked locations in different taxa of the *affinis* subgroup, and we propose a dynamic model of sex chromosome formation and turnover in the *obscura* species group. Our results show that Y genes can find unique paths to escape an unfavorable genomic environment.

## Introduction

Sex chromosomes have formed independently many times from a pair of ordinary autosomes by acquiring a sex-determining gene [1]. In some species groups, such as many fish or reptiles, the proto-X and proto-Y keep recombining over most of their length and evolve little differentiation beyond the sex-determining gene (homomorphic sex chromosomes) [2,3]. However, once the proto-sex chromosomes stop recombining over part or all of their length, they follow different evolutionary trajectories and differentiate genetically and morphologically [4,5]. Ancestral Y chromosomes often are characterized by a loss of most of their ancestral genes, an acquisition of male-specific genes, and an accumulation of repeats and heterochromatin. X chromosomes, in contrast, often evolve dosage compensation [4].

Sex chromosome turnover can be frequent in some groups, especially if the X and Y show little differentiation, but is thought to be rare for heteromorphic sex chromosomes [6]. The highly specialized gene content of old sex chromosomes (i.e. male-fertility genes on the Y) and chromosome-wide regulatory mechanisms (dosage compensation of the X, heterochromatin formation of the Y) is thought to make reversals of highly differentiated sex chromosomes into autosomes increasingly difficult [6]. Recent genomic studies, however, have uncovered turnover of heteromorphic sex chromosomes in multiple taxa. For example, the identity of the X chromosome was found to have changed multiple times across Diptera clades [7].

The evolutionary steps converting an autosome to a sex chromosome have been carefully studied at the molecular level in Drosophila using neo-sex chromosomes [8,9]. The fusion of autosomes to either or both of the ancestral sex chromosomes has repeatedly and independently created neo-sex chromosomes (that is, an X-autosome fusion creates a neo-X, and a Y-autosome fusion creates a neo-Y). Neo-X chromosomes have evolved dosage compensation in multiple Drosophila species [10-13], while neo-Y’s lose most of their ancestral genes, accumulate repetitive DNA, and become heterochromatic [9,14-16].

Genomic comparisons, however, have also started to uncover examples in the reverse direction. In particular, the dot chromosome in Drosophila, a tiny autosome with strongly suppressed recombination, was ancestrally an X chromosome in flies [17]. Indeed, multiple unusual features of this autosome can be better understood in light of its evolutionary history, such as the presence of a dosage-compensation machinery on the dot, or its peculiar expression patterns [18,19]. Intriguingly, comparative analysis of Y-linked genes across Drosophila species also uncovered a Y to autosome reversion in members of the *obscura* species group (the *affinis* and *pseudoobscura* subgroups; see **Figure 1**).

**Figure 1.**
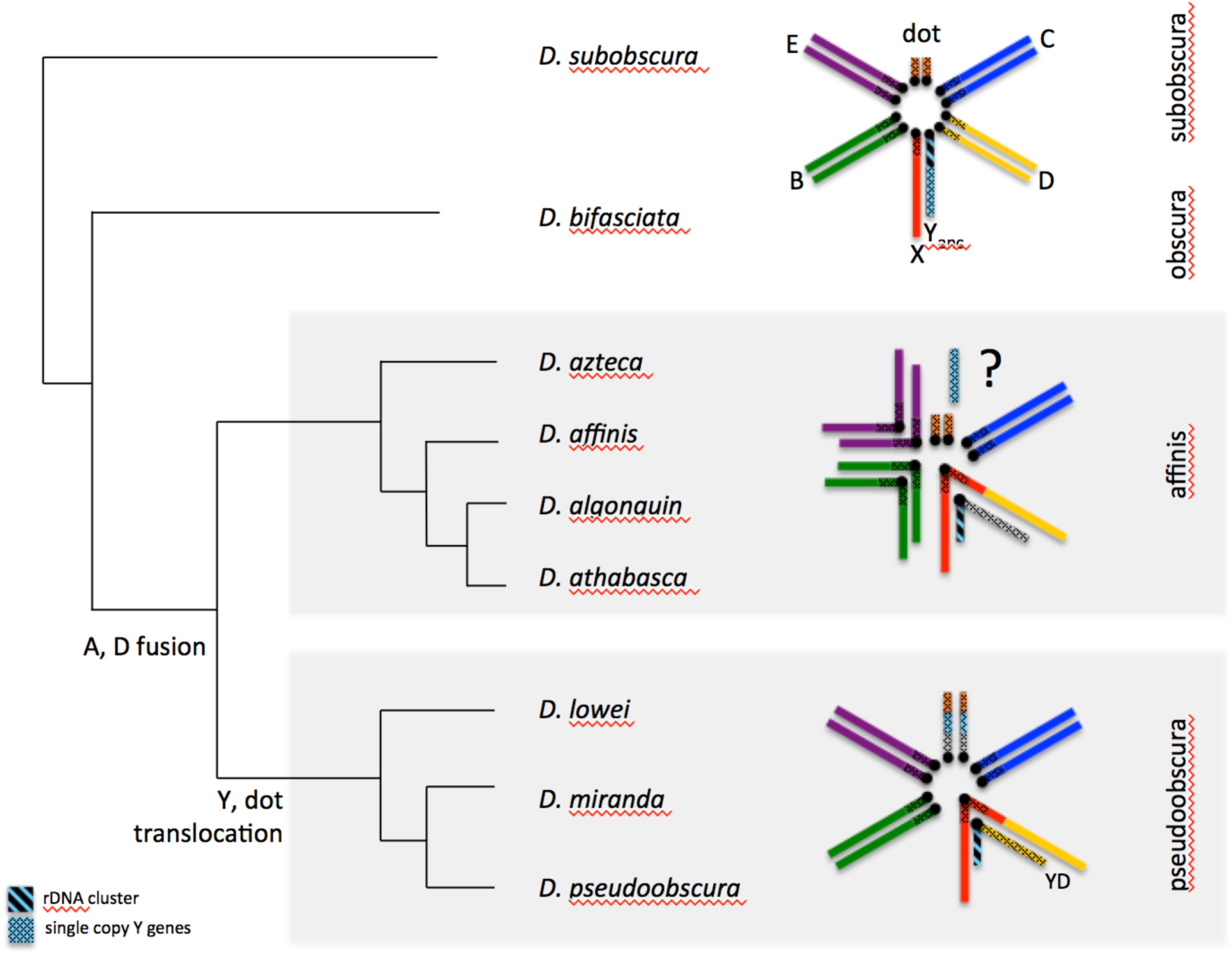
Phylogenetic relationships and karyotype evolution of species in the *obscura* group (male karyotype). Only representative karyotypes that involve transitions of sex chromosomes are drawn (*D. subobscura, D. athabasca, D. pseudoobscura*). The ancestral Y chromosome contains the repetitive rDNA cluster, and single-copy ancestral genes.

In particular, five genes (*Ary, kl-2, kl-3, Ory, Ppr-Y*) that have been shown to be ancestrally present on the Y chromosome of Drosophila were found to be all autosomal in several members of the *affinis* and *pseudoobscura* subgroups [20,21]. Detailed follow-up investigation and genomic analysis showed that the ancestral Y genes are incorporated in the dot chromosome in one piece in both *D. pseudoobscura* and its relative *D. miranda*, suggesting a chromosomal fusion or translocation creating this reversion [16,22,23]. Interestingly, members of the *affinis* and *pseudoobscura* subgroups also share a neo-X chromosome [24,25]. In an ancestor of these lineages, a former autosome (termed Muller element D) fused to the ancestral X chromosome (Muller element A) about 15 MY ago, and the neo-X has evolved the typical properties of an X [12,26](**Figure 1**). The fate of its former homolog (the Muller D element in males not fused to the X) was less clear. In some Drosophila species (such as *D. americana*), X-autosome fusions result in two Y chromosomes (with the unfused chromosome forming a neo-Y), while in others (such as *D. albomicans* and *D. busckii*), the autosomes fuse to both the ancestral X and Y. Males in the *affinis* and *pseudoobscura* subgroups have a single Y chromosome, so it was initially assumed that an unfused neo-Y either completely degenerated, or that the neo-Y became incorporated into the ancestral Y and lost the majority of its genes [24,25].

The discovery of a fusion or translocation between the ancestral Y and the dot chromosome led to an alternative hypothesis about the evolution of the Y in that species [20]. Namely, it was suggested that the ancestral Y and neo-Y did not fuse after the X-autosome fusion, but that putative problems in meiosis that require pairing of multiple sex chromosomes where avoided by the fusion of the ancestral Y with the dot chromosome, and the current Y is a degenerate remnant of the neo-Y of this clade. Support for this notion came from genomic analysis of gene content of the Y chromosome in *D. pseudoobscura*, which was found to be enriched for genes from Muller element D (as would be expected if this chromosome formed from the neo-Y) [27].

Recent work invoking more species, however, hints towards an even more complicated evolutionary history of the sex chromosomes in this clade [28]. In particular, while PCR analysis of the 5 ancestral Y genes confirms their presence in both males and females in most species of the *pseudoobscura* and *affinis* clade, some of them were found to be Y-linked in two species of the *affinis* subgroup: *Ary, kl-2* and *Ory* could only be PCR-amplified from males in *D. athabasca* and *Ary* and *kl-2* showed male-limited PCR-amplification in *D. algonquin* [28]. This was interpreted as the reappearance of Y-linkage for some ancestral Y genes [28]. Here, we use genome analysis to reconstruct the evolutionary history of ancestral Y genes in the *obscura* subgroup (**Figure 1**), taking advantage of chromosome-level assemblies for nine different species (or semi-species) of *D. obscura* flies. Contrary to current belief, we show that the Y-dot fusion/translocation only happened in members of the *pseudoobscura* clade. Surprisingly, we find that ancestral Y genes independently moved away from the Y chromosome to different locations on the autosomes or the X in different species of the *affinis* subgroup. This strongly suggests that Y-linkage of some ancestral Y genes in *D. athabasca* and *D. algonquin* is in fact the ancestral configuration. We propose that translocation of ancestral Y genes can be understood as them escaping from the hostile genomic environment of a neo-Y chromosome, where they suffered the deleterious effects of genetic linkage to a large number of selective targets.

## Results

### Y-dot translocation is only present in the *pseudoobscura* subgroup

The *pseudoobscura* subgroup consists of five described species, and we recently completed chromosome-level genome sequences for three of them [29,16]. For each of the three species (*D. lowei, D. miranda, D. pseudoobscura*), the dot chromosome was assembled in a single contig (**Figure 2, Table 1, Supplemental Figure 1**). Importantly, in each species we detect the five ancestral Y genes assembled in a single genomic fragment, ranging from 180 – 357kb. This fragment is in the same position at the end of each assembled chromosome although inverted in *D. miranda* relative to *D. lowei* and *D. pseudoobscura* (**Figure 2, Supplemental Figure 2)**. Thus, our analysis clearly supports that ancestral single-copy Y genes fused as a single segment to the dot chromosome in flies of the *pseudoobscura* subgroup [16,22,23].

**Table 1.**
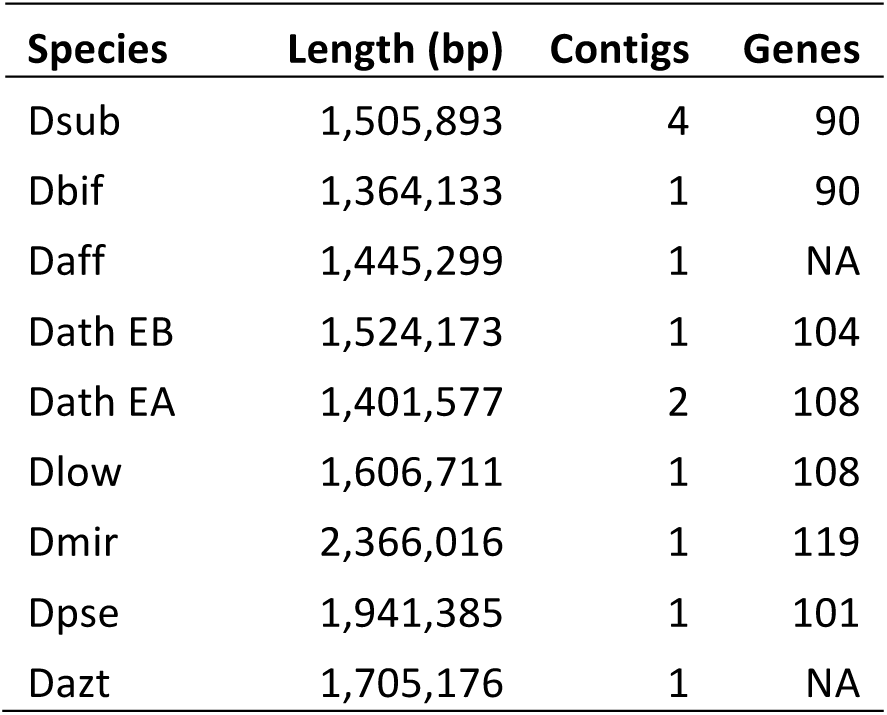
Genome assemblies of the dot chromosome (Muller element F).

**Figure 2.**
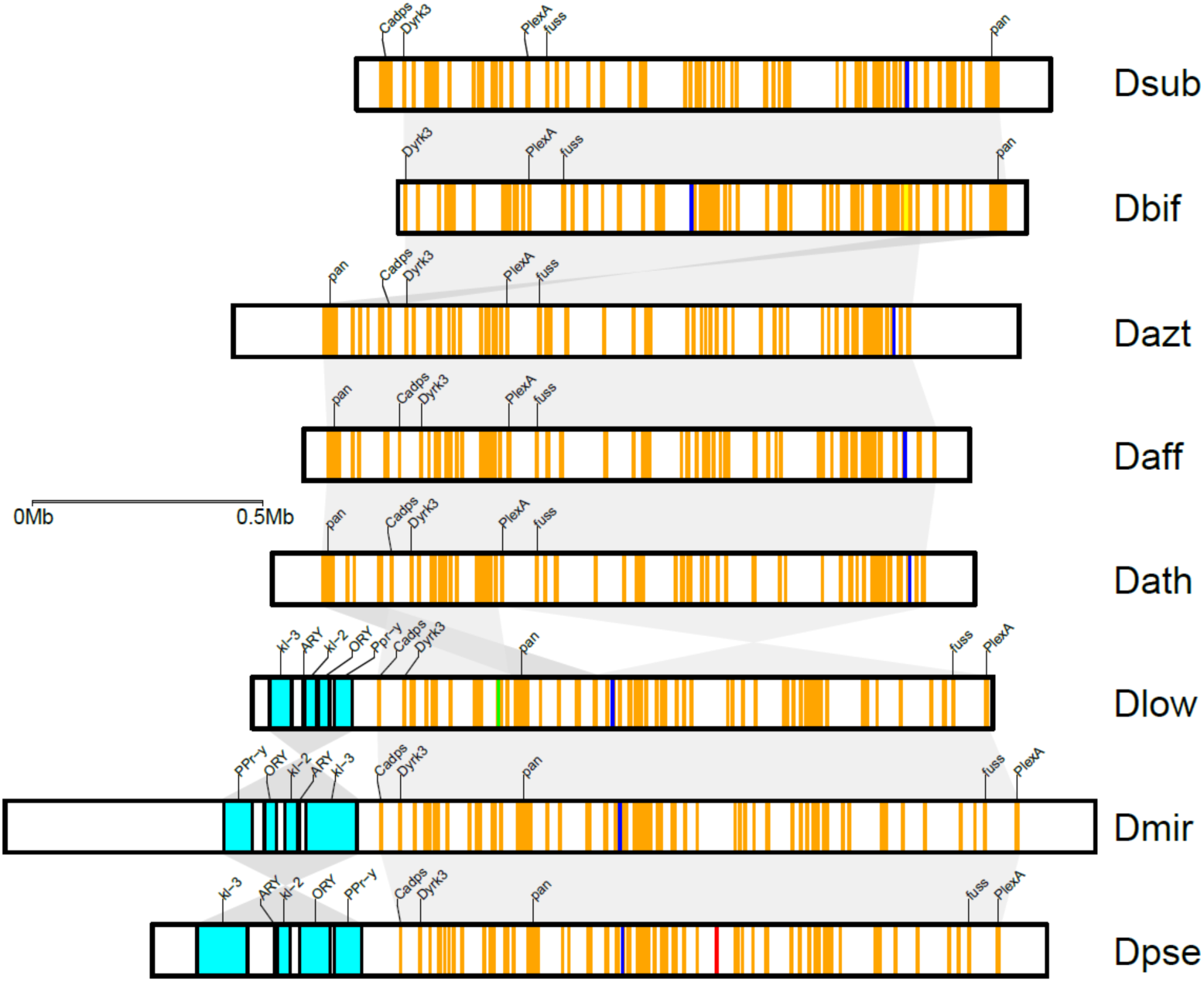
Gene content of the dot chromosome in *obscura* group flies. Shown is the origin of dot genes (orange = Muller F; turquoise = ancestral Y; red = Muller A; green = Muller B; blue = Muller C; yellow = Muller D). Flies from the *pseudoobscura* subgroup all contain ancestral Y genes on the dot chromosome (turquoise), which are absent in other *obscura* group flies, including species from the *affinis* subgroup. The location of best BLAST hit is shown along with the inferred full-length coordinates for ancestral Y genes. Syntenic blocks (> 100kb) shown in grey. Select genes shown overtop each dot chromosome assembly (see Supplemental Figure 1 for all genes).

As expected, we find no ancestral Y genes on the dot chromosomes in *obscura* species that lack the Muller A/D fusion (that is, *D. subobscura* or *D. bifasciata*; **Figure 2, Table 1)**. This is consistent with the hypothesis that the formation of the neo-sex chromosomes causes problems in meiosis, thus driving the fusion or translocation of the ancestral Y chromosome and the dot. Surprisingly, however, we also could not find any ancestral Y genes on the dot chromosome in our high-quality assemblies of two semispecies of *D. athabasca* (Eastern-A (EA) and Eastern-B (EB)), or in a chromosome-level assembly of *D. affinis* or *D. azteca* (**Figure 2, Table 1**). This is unexpected, since the Y-dot translocation is thought to be shared by members of the *affinis* and *pseudoobscura* subgroups. As mentioned above, none of the ancestral Y genes were found to be male-limited in *D. affinis* and most other species in this subgroup, and Y-linkage of *Ary, kl-2* and *Ory* in some lineages was interpreted as these genes gaining Y-linkage secondarily [28].

Consistent with the PCR results [28], we find all five ancestral Y genes in female Illumina libraries from *D. affinis* and *D. azteca* (**Figure 3**). Likewise, we detect *kl-3, Ory* and *Ppr-Y* in reads from a female *D. algonquin* library but not *kl-2* or *Ary*. We find that *kl-3* and *Ppr-Y* are present in female *D. athabasca* but not *Ary, kl-2*, and *Ory* (**Figure 3**). Each of the ancestral Y genes, however, is clearly present in reads from a male genomic library of *D. athabasca*, implying that copies of these genes are found on the male-limited Y chromosome. Genomic read coverage suggests that some of the ancestral Y genes may be present in multiple copies. For example, median read coverage in male and female *D. athabasca* supports one autosomal copy of *kl-3*, one Y-linked copy of *Ary*, while increased male read coverage suggests two Y-linked copies of *kl-2*, and multiple Y-linked copies (or parts of) for *Ory* and *Ppr-Y* (**Figure 3**). Likewise, read-coverage analysis supports multiple (possibly partial) copies of *Ary* and *kl-2* in female *D. azteca*, and possibly multiple (partial) copies of *Ory* and *Ppr-y* in female *D. affinis* (**Figure 3**).

**Figure 3.**
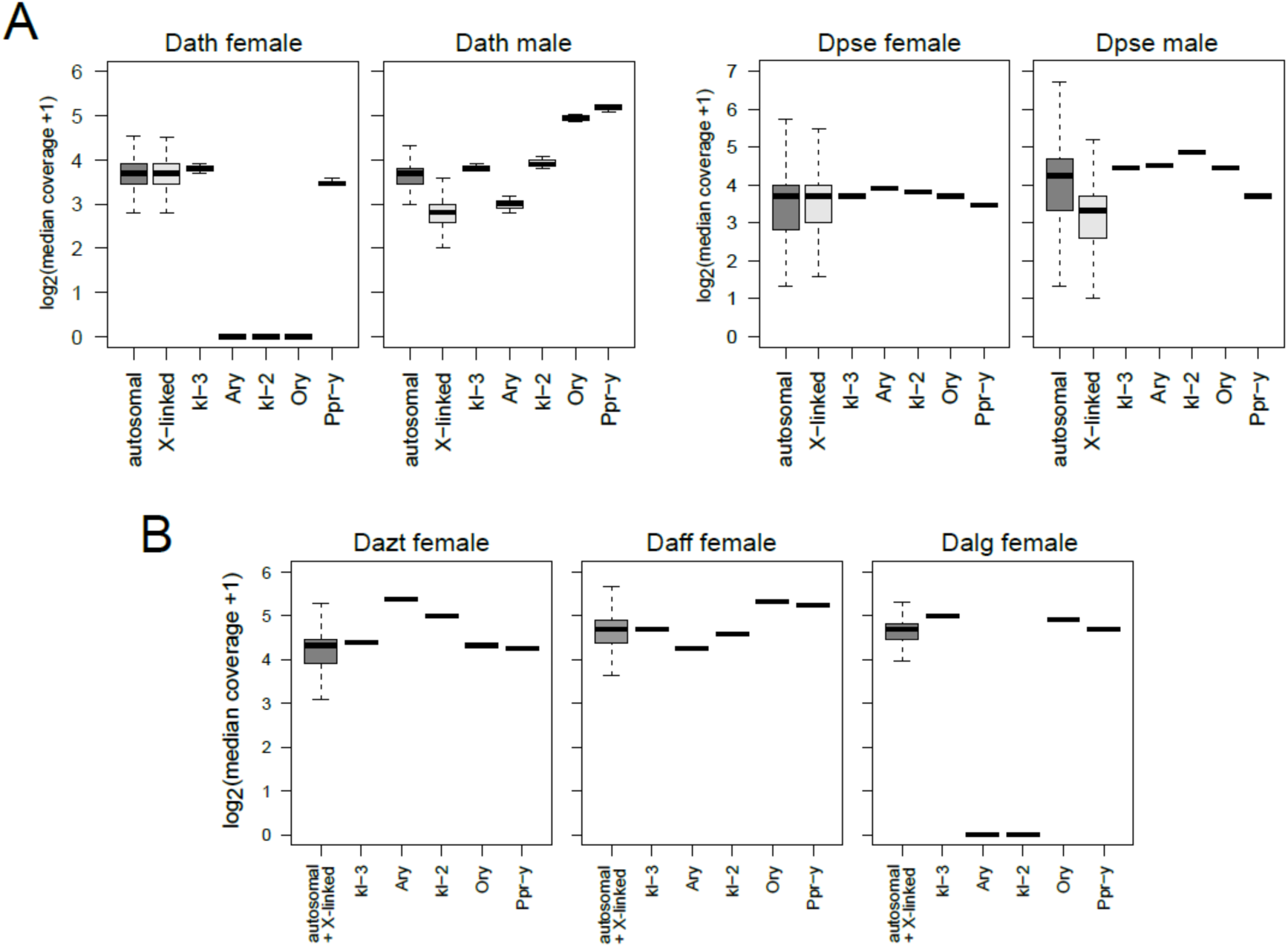
Sex-linkage of ancestral Y genes in *affinis* group flies. A) Shown is sequencing coverage of males and females for genes in *D. athabasca* and *D. pseudoobscura*. B) Shown is genomic coverage of genes for females in *D. azteca, D. affinis* and *D. algonquin*. Outliers not shown for X-linked and autosomal genes.

### Independent incorporation of *kl-3* and *Ppr-Y* on Muller B of *D. athabasca*

If not on the dot chromosome, where are ancestral Y genes found in *affinis* flies? Consistent with our coverage analysis and PCR results [28], we find *kl-3* and *Ppr-Y* to be contained in both of our female assemblies of EA and EB *D. athabasca*, but not *Ary, kl-2* and *Ory*. Surprisingly, however, both of these genes are located on Muller B, in different chromosomal locations (**Figure 4, Supplemental Table 1**). In particular, we find *Ppr-Y* on the short arm of Muller B (around 1.7 Mb), while *kl-3* is located on the long arm (around 37.7Mb) in the EB assembly, and their locations are conserved in the EA semispecies. Thus, unlike the Y- to dot translocation in the *pseudoobscura* subgroup, we find that *kl-3* and *Ppr-Y* moved independently away from the Y chromosome to a different autosome in *D. athabasca*. We could not find *Ary, kl-2* and *Ory* in our female assembly by BLAST, consistent with our Illumina read mapping and PCR results [28].

**Figure 4.**
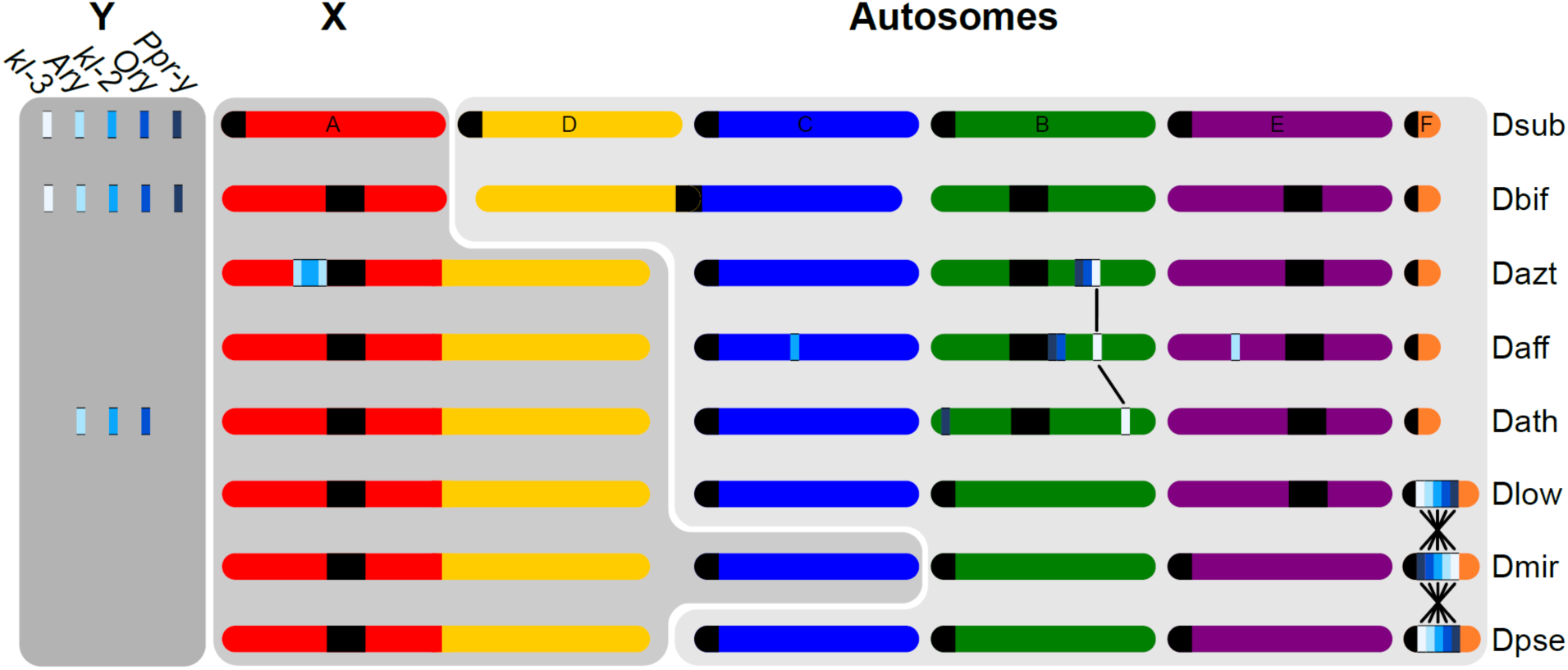
Schematic representation of location of ancestral Y genes in *obscura* group flies. Shown is the approximate genomic location of the five Y_anc_ genes based on high-quality genome assemblies. The presence/absence of Y_anc_ genes on the Y chromosome is inferred from genomic coverage patterns. Muller elements are color-coded as in Figure 1 and identified in *D. subobscura*. The vertical lines connect genes found in homologous positions. Note that Muller C fuses to Muller A-AD in some *D. athabasca*, but for simplicity is not shown.

Ancestral Y genes in *D. melanogaster* are expressed almost exclusively in testis [30]. Testis expression patterns of ancestral Y genes have been conserved for *pseudoobscura* subgroup flies, where they moved as a single piece to the dot chromosome [16,23]. We used RNA-seq data from different male and female samples (male and female whole larvae, male and female adult and larvae heads; adult testis and ovaries) to investigate sex- and tissue-specific expression patterns of ancestral Y genes from both EA and EB *D. athabasca.* Consistent with these genes having important functions in Drosophila spermatogenesis, we find that they are all highly expressed in testis of *D. athabasca* (**Table 2**). Thus, both the genes that have stayed behind on the Y chromosome (*Ary, kl-2, Ory*) but also those that moved to an autosome (*Ppr-Y, kl-3*) have maintained their male-specific expression profile.

**Table 2.** Gene expression of ancestral Y genes from different tissues and sexes of two *D. athabasca* semispecies (Eastern-A and Eastern-B). Values are in FPKM (fragments per kilobase of transcript per million mapped reads).

|  |  | *kl-3* | *Ary* | *kl-2* | *Ory* | *Ppr-y* |
| --- | --- | --- | --- | --- | --- | --- |
| *Eastern-B* |  |  |  |  |  |  |
| male | whole larvae | 0.8 | 0 | 0 | 0 | **1.3** |
| male | larval heads | 0 | 0 | 0 | 0 | 0 |
| male | testes | **67.2** | 0 | **11.3** | **39.54** | **73.8** |
| male | heads | 0 | 0 | 0 | 0 | 0.7 |
| female | whole larvae | 0.4 | 0 | 0 | 0 | 0 |
| female | larval heads | 0 | 0 | 0 | 0 | 0 |
| female | ovaries | 0 | 0 | 0 | 0 | **8.3** |
| female | adult heads | 0 | 0 | 0 | 0 | 0 |
| *Eastern-A* |  |  |  |  |  |  |
| male | whole larvae | **2.5** | 0 | 0.6 | **2.3** | **3.6** |
| male | larval heads | 0.0 | 0 | 0 | 0 | 0.7 |
| male | testes | **81.3** | **10.2** | **12.1** | **44.5** | **292.2** |
| male | heads | **1.4** | 0 | 0 | 0 | 0 |
| female | whole larvae | 0.6 | 0 | 0 | 0 | 0 |
| female | larval heads | 0 | 0 | 0 | 0 | 0 |
| female | ovaries | 0 | 0 | 0 | 0 | 0 |
| female | adult heads | 0 | 0 | 0 | 0 | 0 |

Thus, our analysis confirms that *Ary, kl-2* and *Ory* are still present in the male genome of *D. athabasca* but not in females, i.e. these genes are located on the Y chromosome in this species. This is consistent with the PCR results of [28]. However, while they assumed that the Y-dot translocation was shared by *pseudoobscura*/*affinis* flies and therefore interpreted their PCR screen of *Ary, kl-2* and *Ory* being only present in males as them becoming Y-linked secondarily, a simpler explanation is that these genes never left the Y. Our data indicate that the Y-dot translocation is unique to flies in the *pseudoobscura* subgroup, and we show that *kl-3* and *Ppr-Y* independently became autosomal in *D. athabasca*.

### Independent Y gene gain in *D. affinis* and *D. azteca*

In most species in the *affinis* subgroup (of which *D. athabasca* is a member), ancestral Drosophila Y genes are present in both sexes (**Figure 3**) [20,28]. This was interpreted as a single Y-dot translocation moving all ancestral Y genes to an autosome [22,28], but a lack of Y genes on the dot of *D. athabasca* and *D. affinis* argues against this scenario, and our results from *D. athabasca* suggest that ancestral Y genes may have been moved independently to autosomal locations in different species. To test this hypothesis, we analyzed high-quality genomes from *D. affinis*, a sister species to *D. athabasca* from which it diverged <3MY ago [31], and *D. azteca* (which diverged <6MY ago; Beckenbach *et al.* 1993), two species for which all ancestral Y genes were found in both sexes. Indeed, we find copies for each ancestral Y gene in the female assembly of both species, but at strikingly diverse genomic locations (**Figure 4**).

In particular, four of the 5 ancestral Y genes are found on different chromosomal locations in the *D. affinis* genome: *kl-2* is on Muller-C (at 9.2Mb), *kl-3* is on Muller B (10.0 Mb), *Ary* is on Muller E (9.8 Mb) and *Ory* and *Ppr-Y* appear to have translocated together onto Muller B (15.6 Mb). Comparisons of flanking regions suggest that the translocation of *kl-3* occurred in an ancestor of *D. affinis* / *D. athabasca*, as *kl-3* is surrounded by the same genes in both species (**Figure 5A**). *Ppr-Y*, on the other hand, is found on non-homologous positions between *D. affinis* / *D. athabasca*, suggesting that this gene moved independently to Muller B in the two species. The *kl-2* translocation on Muller C in *D. affinis* appears to have only occurred in this species (**Figure 5A**).

**Figure 5.**
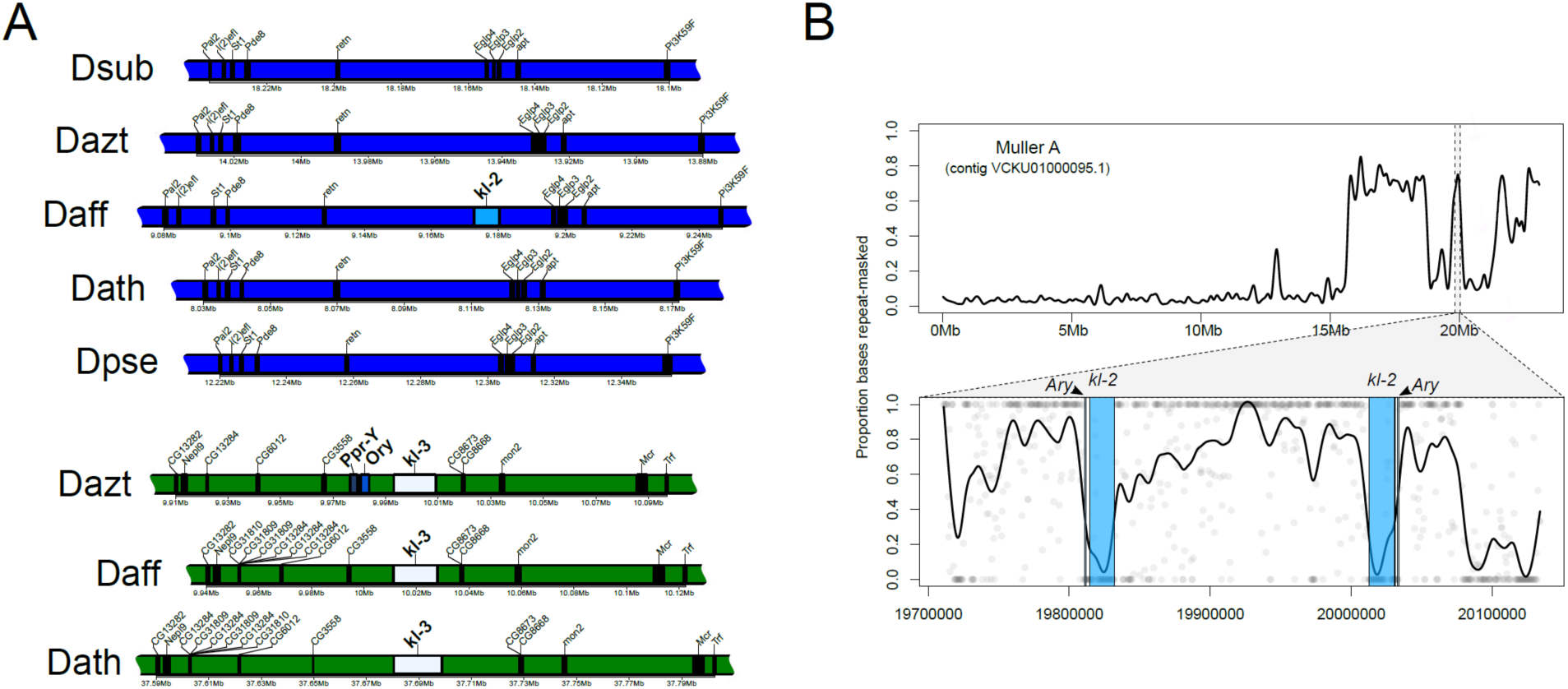
Details of ancestral Y gene translocations. A) Local alignments around *kl-2* indicate that this gene translocated to a region on Muller C (blue) in *D. affinis*. Local alignments of the translocation of *kl-3* on Muller B (green) show it is in a homologous position in *D. azteca, D. affinis*, and *D. athabasca. Ppr-y* and *Ory* appear absent from the region in *D.affinis* and *D. athabasca*. B) *Ary*/*kl-2* are duplicated on XL (Muller A) of *D. azteca*, resembling palindromes found on the human Y chromosome. Shown above is a LOESS smoother fit to the proportion of bases repeat-masked in 500bp windows. Below highlights the genomic interval harboring *Ary/kl-2.* Dots show individual 500bp window estimates with a LOESS smoother fit to the genomic interval.

Likewise, ancestral Y genes in *D. azteca* are located in different regions of the female genome assembly (**Figure 4**). *Ppr-Y, kl-3* and *Ory* are found next to each other on Muller B (10.0 Mb), suggesting that they moved in one piece, and comparisons of flanking genes suggest that *kl-3* is located on a homologous position in *D. affinis* and *D. athabasca* (**Figure 5A**). Comparisons of this region in the *D. pseudoobscura* and *D. subobscura* genomes show that this Y gene translocation occurred at an *affinis* subgroup specific inversion breakpoint (i.e, breakpoint relative to the *subobscura*/*pseudoobscura* subgroups), which limits our understanding of the size of the translocation. Our findings suggests that *kl-3* moved to Muller B in an ancestor of the *affinis* subgroup, and this initial translocation may have also included *Ppr-Y* and *Ory*, which were lost in the lineage leading to *D. athabasca*. An additional inversion may have moved *Ppr-Y* and *Ory* close to the pericentromere in *D. affinis* (but note that the long arm of Muller B appears completely syntenic between *D. affinis* and *D. azteca*, arguing against simple inversions; **Supplemental Figure 3**). *Ppr-Y* and *Ory* could also have moved secondarily onto the long arm of Muller B in *D. azteca* and independently in *D. affinis*, and *Ppr-Y* moved independently onto the short arm of Muller B in *D. athabasca.* Under either scenario, our results support a dynamic evolutionary history of ancestral Y gene movement in flies of the *affinis* subgroup. In *D. azteca*, we find that *Ary* and *kl-2* moved together to Muller A (the ancestral X chromosome), and both appear to be duplicated next to each other in opposite directions, with about 180 kb of sequence in between them (**Figure 5B**). This insertion appears close to, or in, the pericentromere as the region has high repeat density and shows sequence similarity with pericentromeric regions in *D. athabasca* and *D. affinis*. (**Figure 5B**). The sequence in between the *Ary*/*kl-2* duplication is almost entirely composed of repeats (75.1% repeat masked), and may thus be derived from the Y chromosome. The overall arrangement of *Ary* and *kl-2* resembles the palindrome structure of multicopy genes on the human Y chromosome [32,33], but it is unclear if this arrangement arose before or after these genes moved onto Muller A.

In summary, the absence of the Y-dot fusion, and a lack of conservation of location for most ancestral Y genes in the *affinis* subgroup indicates that genes moved away independently from the Y in this clade. Y genes in *D. melanogaster* can be gigantic, due to huge introns [30] and require unique gene expression programs [34]. Multiple independent translocations of ancestral Y genes suggest that the Y chromosome may have been smaller in *obscura* subgroup flies compared to *D. melanogaster*, which is consistent with karyotypic findings [23].

## Discussion

The *obscura* species group of Drosophila provides a fascinating clade to study karyotype evolution [29], and it contains multiple sex chromosome transitions. Neo-sex chromosomes formed independently in different clades, including the fusion of the ancestral X with Muller D roughly 15MY ago, but also more recent fusions of Muller C with the Y chromosome in *D. miranda* and in some semispecies of *D. athabasca*, which allows us to reconstruct the events transforming an autosome into differentiated sex chromosomes. Intriguingly, however, we also observe the independent incorporation of ancestral Y genes in different species of *affinis* and *pseudoobscura* subgroup flies.

The ancestral Y of Drosophila contains both single-copy genes and the multicopy rDNA cluster [22,35,36]. FISH studies have shown that the rDNA cluster is present on both the X and Y chromosome in multiple species of *obscura* flies, including members from the *obscura, affinis* and *pseudoobscura* subgroups [22]. This suggests that this is the ancestral configuration of the rDNA cluster, and its location on the Y was maintained even in species where single-copy Y genes translocated to the dot (*pseudoobscura* subgroup) or other chromosomes (*affinis* subgroup).

While we cannot reconstruct the early events of sex chromosome evolution in the *obscura* group with certainty, we propose the following model that accounts for the genomic location of ancestral and newly formed sex-linked genes (**Figure 6**). In an ancestor of the *affinis*/*pseudoobscura* subgroups, the ancestral X fused to Muller D, and formed the second arm of the X chromosome found in all species belonging to these two subgroups. Such a fusion leaves the unfused Muller D, and the ancestral Y chromosome, and their fate has been less clear. Given Y-linkage of rDNA genes in species from all groups in *obscura* flies, this suggests that the rDNA cluster was ancestrally on the Y, and all species have incorporated at least part of the ancestral Y into their current Y [22]. Additionally, some species in the *affinis* subgroup (*D. athabasca, D. algonquin*) have maintained ancestral single-copy Y genes on their current Y (see above; [28]. Furthermore, an over-abundance of Muller D genes was found on the current Y chromosome of *D. pseudoobscura* and *D. miranda* [16,20,27], suggesting that Muller D (or part of it) also became incorporated into the Y of *pseudoobscura* subgroup flies. Thus, the simplest explanation for the current gene content of the Y in species with the X-D fusion is that Muller D also fused to the ancestral Y. Indeed, it is possible that the Y-D fusion actually preceded the X-D fusion, mimicking the current Y-autosome fusions found in *D. miranda* and *D. athabasca*, which would leave males with two un-linked X chromosomes. The fusion between either the X or Y chromosome and Muller D would generate a trivalent in males (that is, an X-D fusion creates two Y chromosomes in males, while a Y-D fusion would create two X’s in males that need to pair with one Y) and create problems in meiosis, resulting in higher rates of aneuploidy. This could rapidly select for a second fusion of Muller D with the unfused sex chromosome, as was experimentally demonstrated in a hybrid population of *D. albomicans* (a species that contains both a X-autosome and Y-autosome fusion) and its sister species *D. nasuta* that lacks neo-sex chromosomes [37]. If Muller D fused with both the ancestral X and Y, this should alleviate problems associated with segregating a trivalent. Ancestral Y genes then secondarily translocated to autosomal or X-linked locations, either as a single unit to the dot chromosome in an ancestor of the *pseudoobscura* subgroup, or individually in species of the *affinis* subgroup (**Figure 6**).

**Figure 6.**
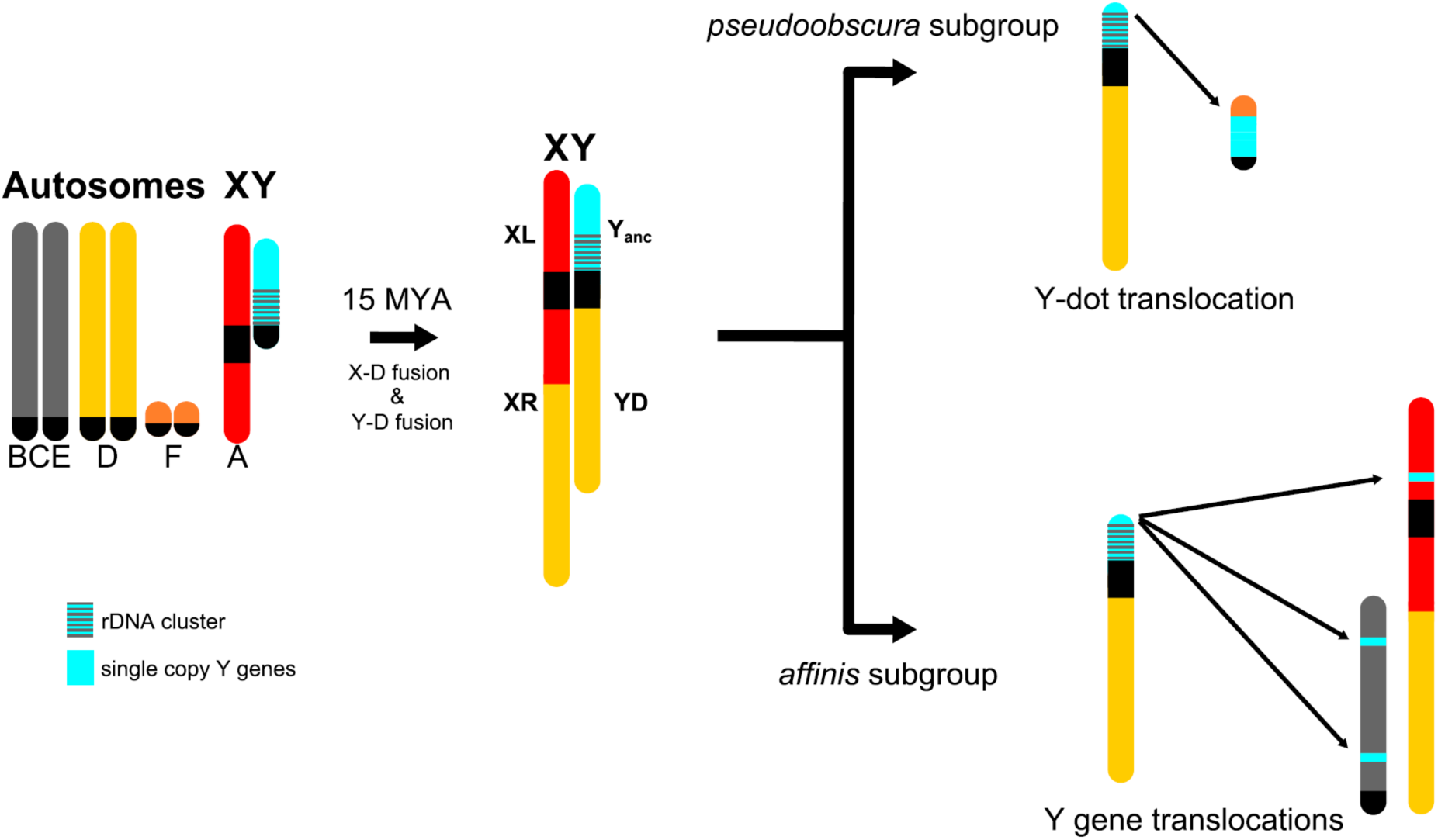
Model of sex chromosome evolution in the *obscura* group. In an ancestor of the *affinis* and *pseudoobscura* subgroups, the ancestral X (Muller A) and Muller D fused ∼15 million years ago. We hypothesize that the ancestral Y, which carries the rDNA cluster and single-copy Y_anc_ genes, also fused to Muller D, which would explain Y-linkage of the rDNA cluster in all species, and Y-linkage of Y_anc_ genes in several species. In the *pseudoobscura* subgroup, single-copy Y_anc_ genes translocated in one fragment to the dot chromosome, leaving behind (fragments of) Y_anc_ genes on the Y chromosome. In the *affinis* group, Y_anc_ genes moved independently to different autosomal and X-linked location in different clades/species.

What might drive the relocation of ancestral Y genes? Becoming linked to a gene-rich chromosome will present a novel challenge for genes with important functions in spermatogenesis that have managed to survive for millions of years on a non-recombining Y chromosome. In particular, evolutionary models to explain the degeneration of a Y are based on interference among selected mutations on a non-recombining chromosome [38,39]. Theory and computer simulations have shown that the magnitude of selection interference, and thus the rate of degeneration, depends on the number of functional genes present on the Y chromosome [40]. Gene loss is highest on a gene rich Y chromosome, but declines rapidly as active genes are lost [40]. While old, degenerate Y chromosomes may provide safe havens for important male-specific genes, ancestral Y genes suffer the deleterious effects of genetic linkage to more selective targets when fused to an autosome containing 1000s of functional genes. Their translocation may thus be driven to avoid mutation accumulation and degeneration on the neo-Y where purifying selection is highly impaired. This resembles the fate of a Y gene (*kl-5*) in the *testacea* group species of *Drosophila* that duplicated to the dot chromosome [41]. The dot, like the Y chromosome, lacks recombination but contains about seven times more genes. It was shown that slightly deleterious mutations have accumulated in the dot-linked copy of *kl-5* faster than in the Y-linked copy [41], consistent with the copy on the dot suffering the deleterious effects of genetic linkage to more selective targets compared with the Y chromosome.

Thus, our findings suggest a turbulent history of Y genes in the *obscura* group. After being protected from the accumulation of deleterious mutations on the gene-poor ancestral Y for millions of years, linkage to Muller D would have caused massive selective interference and degeneration of these genes. Y genes in the *pseudoobscura* subgroup escaped to a suboptimal genomic environment on the dot chromosome, while ancestral Y genes in the *affinis* subgroup began to duplicate or translocate to other autosomal locations. Therefore, a highly degenerate Y chromosome may not be that inhospitable as commonly assumed, and may instead be a safe haven for male-beneficial genes.

A noticeable commonality between several of the ancestral Y gene translocations is that their autosomal copies are often found near heterochromatin. Ancestral Y genes fused to the heterochromatic dot chromosome in the *pseudoobscura* subgroup, *Ary*/*kl-2* are adjacent the pericentromere on Muller A in *D. azteca*, and *Ory*/*Ppr-Y* are near the pericentromere on Muller B in *D. affinis* (**Figure 4**). In addition, we found fragments of Y-linked genes in the pericentromeres of several other species (**Supplemental Table 2**). This suggests that ancestral Y genes may have an affinity for heterochromatin, and non-allelic homologous recombination between the repeat-rich Y chromosome and repetitive autosomal regions could facilitate movement of ancestral Y genes. Additionally, heterochromatin may be a preferential location for ancestral Y genes, as their regulatory machinery has evolved in a heterochromatic environment on the ancestral Y.

## Methods

Seven of the *Drosophila obscura* group genome assemblies (*D. athabasca* EA and EB, *D. lowei, D. miranda, D. pseudoobscura, D. subobscura, D. bifasciata*) used in our analyses are described in detail in [16], [29] and Bracewell et al. (submitted) and are available through GenBank. For *D. affinis*, we used a newly generated PacBio-based genome assembly kindly provided by Rob Unckless. For *D. azteca*, we downloaded the most recent version from GenBank (accession GCA_005876895.1) and additional details can be found at NCBI Bioproject PRJNA475270. To assign *D. azteca* contigs/scaffolds to Muller elements we used D-Genies [42] to perform whole genome alignments with our other chromosome-level genome assemblies. During genome alignments and BLAST searches (below), we flagged contig VCKU01000055.1 as chimeric as it is a composite of sequences that map uniquely to different pericentromeric regions on all chromosomes in other assemblies. After identifying the Muller F from all assemblies, we generated alignments and dot plots using MUMmer [43] with NUCmer --mum -c 200 and mummerplot with the --filter option.

To find ancestral Y genes we used the annotation file (gtf) and dot (Muller F) assembly from [23] along with gffread (https://github.com/gpertea/gffread) to generate transcripts of ancestral-Y genes for use in BLASTN searches with *obscura* group genome assemblies (above). We retained the longest transcript for these five genes (see Supplemental fasta file). To further confirm our BLASTN results, we downloaded all *D. melanogaster* translations (r6.30) from FlyBase (flybase.org) and used TBLASTN to again search all obscura group assemblies. BLASTN and TBLASTN searches had colocalized hits, except for *Ppr-Y*, which was only found using BLASTN searches with the *obscura* group transcript. Results from BLASTN searches can be found in Supplemental Table 2. Only hits with ≥ 80% sequence identify were kept. BLAST Searches of *D.azteca* for *kl-3, Ppr-Y* and *Ory* also returned high-scoring hits to contig VCKU01000055.1 which are not shown due to it likely being an assembly artifact.

To estimate sequencing coverage over genes we generated whole genome sequencing data (Illumina) for an individual female of *D. azteca and D. affinis*, and males and females of *D. athabasca.* We extracted DNA from each individual using a Qiagen DNeasy kit following manufacturers recommendations. DNA libraries were prepared using the Illumina TruSeq Nano Prep kit and sequenced on a Hiseq 4000 with 100 bp PE reads. We downloaded *D. algonquin* Illumina data that have previously been deposited with the SRA (accession SRR5768634). To estimate coverage over genes, we used as a reference the longest *D. athabasaca* (EB) transcript for each gene from MAKER annotations [29] along with the *D. pseudobscura* transcripts for *kl-2, Ary* and *Or*y. We then used BWA MEM [44] to map all paired-end Illumina reads as single-end reads to these transcripts. Samtools [45] was used to manipulate files and coverage over each transcript (gene) was estimated from the bam files using bedtools genomecov and groupBy [46].

We characterized gene expression of the five ancestral Y genes by analyzing RNA-seq data from [29]. We first cleaned raw Illumina reads using SeqyClean (https://github.com/ibest/seqyclean) and then used the HiSat2 [47], Samtools [45] and the StringTie [48] pipeline to estimate FPKM for all expressed transcripts.

Plots of Muller F assemblies and locations of ancestral Y insertions were created using Karyoploter [49]. Genes shown with *D. melanogaster* gene names are the result from TBLASTN searches (above) and only top hits with ≥ 50% sequence identity were plotted. To estimate repeat density in *D. azteca* we used Repeatmasker version 4.0.7 [50] with the -no_is and -nolow flags and the Repbase *Drosophila* repeat library (downloaded March 22, 2016, from www.girinst.org). The proportion of repeat-masked bases (Ns) in non-overlapping windows along the masked genome was determined using bedtools nuc.

## Data availability

All sequence data generated for this study has been deposited with the NCBI SRA and can be found under the NCBI BioProject PRJNA589338.

---

**Supplemental Figure 1.**
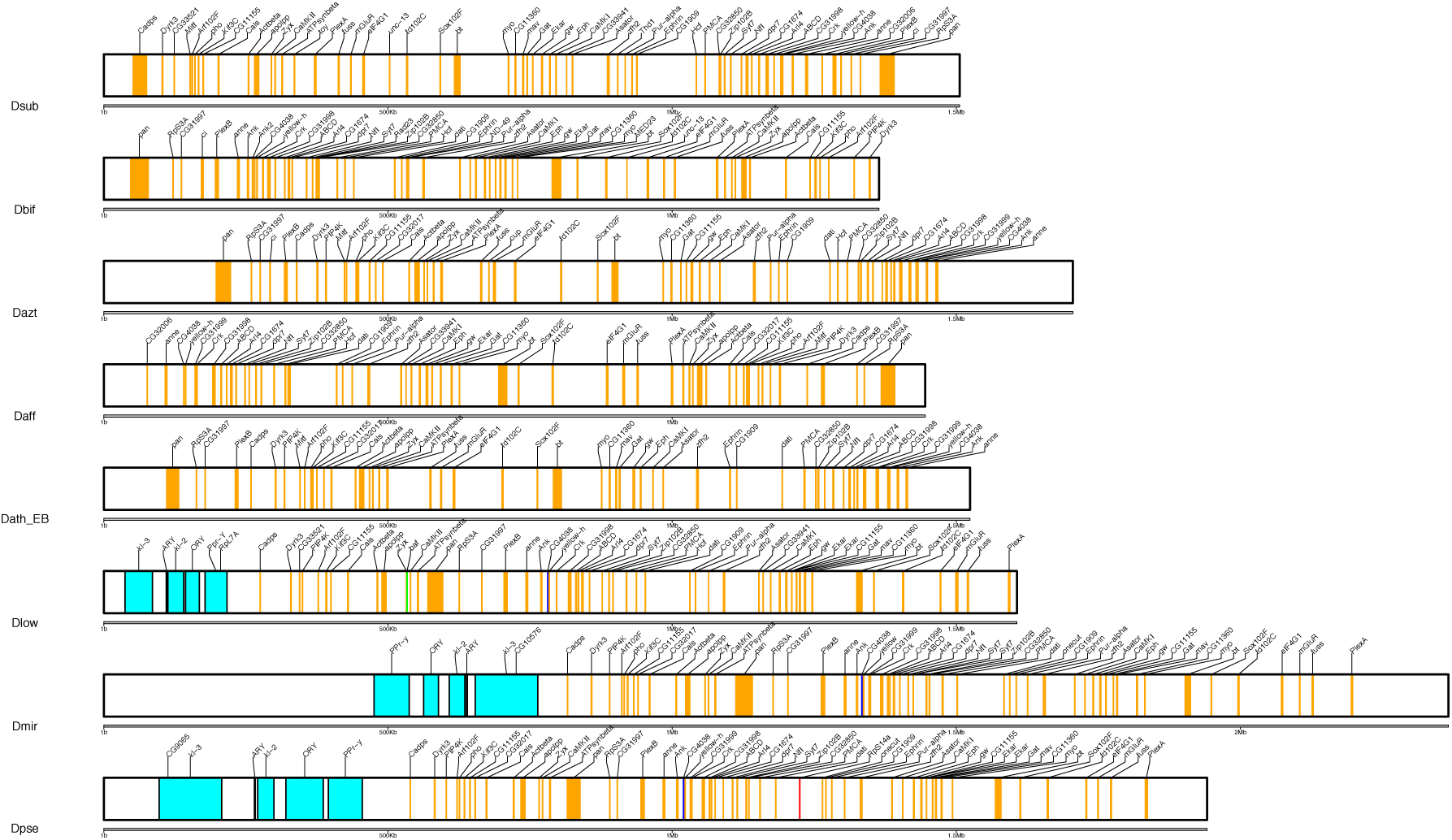
Details of dot chromosomes (Muller F) from eight species in the *Drosophila obscura* group. Drosophila melanogaster (Dmel) gene names shown. Genes are color-coded based on their location in Dmel (orange = Muller F; turquoise = ancestral Y; red = Muller A; green = Muller B; blue = Muller C; yellow = Muller D).

**Supplemental Figure 2.**
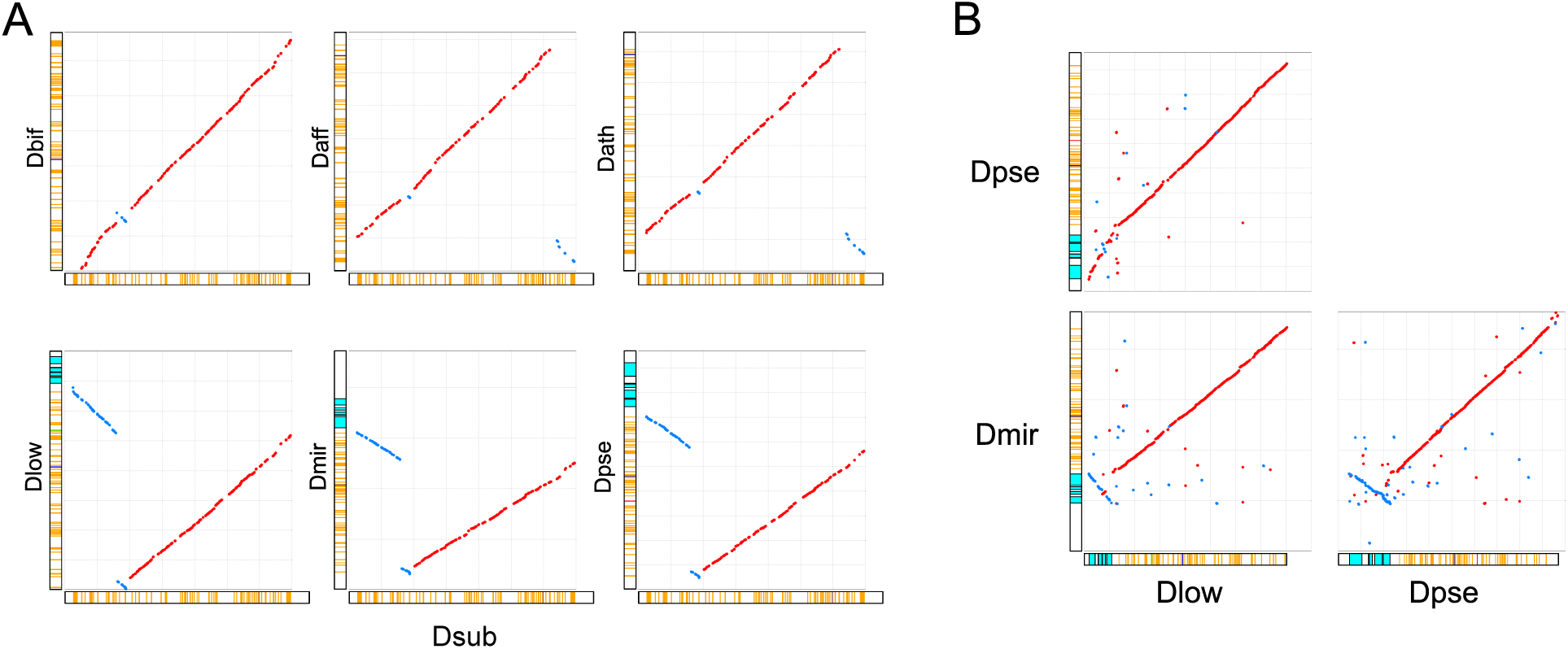
Dot plot comparisons (MUMmer) of the dot chromosome (Muller F) between species. Dot chromosome depictions on the axes are from Figure 2. A) Comparisons of the *Drosophila subobscura* dot (X axis) to *D. bifasciata, D. affinis, D. athabasca* (EB), *D. lowei, D. miranda* and *D. pseudoobscura*. B) All pairwise comparisons of *pseudoobscura* group species harboring the Y-dot fusion.

**Supplemental Figure 3.**
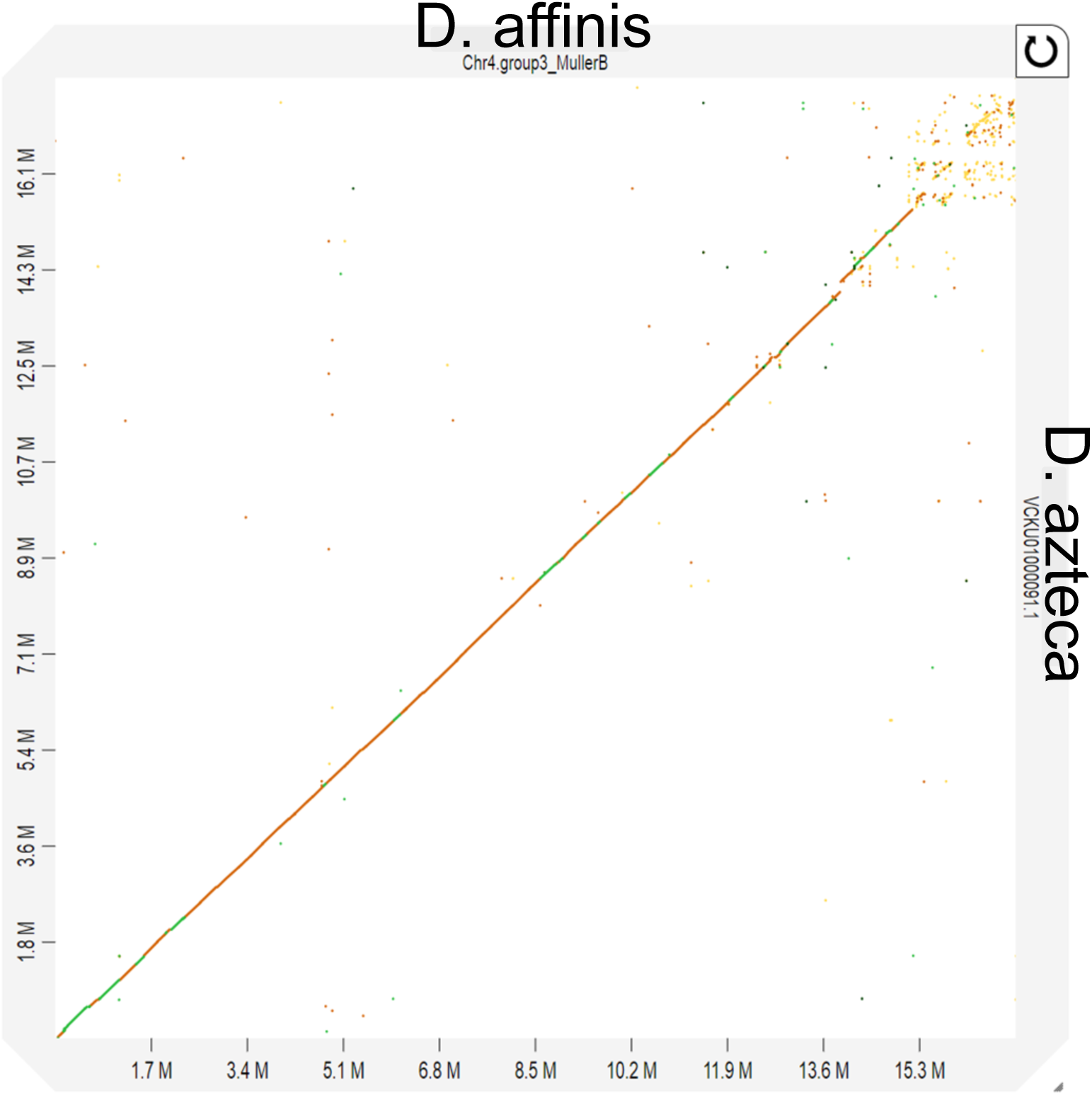
Dot plot comparisons (MUMmer) of Muller B between *D. affinis* and *D. azteca.*

